# Host tissue proteomics reveal insights into the molecular basis of *Schistosoma haematobium*-induced bladder pathology

**DOI:** 10.1101/2021.11.23.469626

**Authors:** Derick N. M. Osakunor, Kenji Ishida, Olivia K. Lamanna, Mario Rossi, Louis Dwomoh, Michael H. Hsieh

**Affiliations:** Division of Urology, Department of Surgery, Sheikh Zayed Institute for Pediatric Surgical Innovation, Children’s National Hospital, Washington, District of Columbia, United States of America; Institute of Molecular, Cell and Systems Biology, University of Glasgow, Glasgow, Scotland, United Kingdom, Departments of Urology, Department of Pediatrics, and Department of Microbiology, Immunology, and Tropical Medicine, School of Medicine and Health Sciences, The George Washington University, Washington, District of Columbia, United States of America

**Author notes:** Department of Applied Clinical and Biotechnology Sciences, University of L’Aquila, L’Aquila, Italy (Mario Rossi).

**Keywords:** Bladder, Bladder wall injection, Mice, Proteomics, *Schistosoma haematobium*, Schistosomiasis, Urogenital schistosomiasis

## Abstract

**Background:** Urogenital schistosomiasis remains a major public health concern worldwide. In response to egg deposition, the host bladder undergoes gross and molecular morphological changes relevant for disease manifestation. However, limited mechanistic studies to date imply that the molecular mechanisms underlying pathology have not been well-defined. We leveraged a mouse model of urogenital schistosomiasis to perform for the first time, proteome profiling of the early molecular events that occur in the bladder after exposure to *S. haematobium* eggs, and to elucidate the protein pathways involved in urogenital schistosomiasis-induced pathology.

**Methods:** Purified *S. haematobium* eggs or control vehicle were microinjected into the bladder walls of mice. Mice were sacrificed seven days post-injection and bladder proteins isolated and processed for proteome profiling using mass spectrometry.

**Results:** We demonstrate that biological processes including carcinogenesis, immune and inflammatory responses, increased protein translation or turnover, oxidative stress responses, reduced cell adhesion and epithelial barrier integrity, and increased glucose metabolism were significantly enriched in *S. haematobium* infection.

**Conclusion:** *S. haematobium* egg deposition in the bladder results in significant changes in proteins and pathways that play a role in pathology. Our findings highlight the potential bladder protein indicators for host-parasite interplay and provide new insights into the complex dynamics of pathology and characteristic bladder tissue changes in urogenital schistosomiasis. The findings will be relevant for development of improved interventions for disease control.

**Author summary:** The molecular mechanisms underlying the urinary and genital pathology from urogenital schistosomiasis have not been well-defined. This has mainly been due to limited mechanistic studies and the lack of a suitable animal model for *S. haematobium* infection. We leveraged a mouse model of urogenital schistosomiasis, along with proteomics analysis, to determine the early molecular events that occur in the bladder after exposure to *S. haematobium* eggs, and to define protein pathways that may be involved in morbidity from urogenital schistosomiasis infection. Our results show that *S. haematobium* egg deposition in the bladder results in significant changes in proteins and pathways that play a role in pathology. Biological processes including carcinogenesis, immune and inflammatory responses, increased protein translation or turnover, oxidative stress responses, reduced cell adhesion and epithelial barrier integrity, and increased glucose metabolism are significantly enriched in *S. haematobium* infection. Our findings highlight important protein indicators of host-parasite interactions in the bladder and their association with tissue changes in urogenital schistosomiasis. The findings presented here will be relevant for development of improved interventions for disease control, and contribute to realizing the 2021-2030 NTD goals by demonstrating evidence on the early events that occur during schistosome infection.

## Introduction

Second only to malaria among the most devastating parasitic diseases with the greatest economic and public health impact, schistosomiasis affects over 250 million people worldwide and accounts for the loss of over 2.5 million disability-adjusted life years (DALYs) [1–3]. It is caused by trematodes of the *Schistosoma* genus; the most common species, *Schistosoma haematobium,* causes urogenital schistosomiasis, which accounts for about two-thirds of all schistosomiasis cases [4]. Adult worm pairs typically reside in the venous plexus of the bladder and lay eggs in bladder tissue, which are excreted and can be detected in the urine [4]. Although the nature and magnitude of disease may vary, most morbidity and mortality are mainly due to host immunological reactions to eggs deposited within the walls of the urinary tract. This can lead to increased urinary tract infections, hematuria and proteinuria due to a compromised urothelium [5, 6], inflammatory fibrosis of the bladder, bladder carcinogenesis, and obstructive renal failure [7, 8]. Schistosomiasis-associated bladder cancer is a frequent and dire consequence of urogenital schistosomiasis, with incidence rates estimated at about 3-4 cases per 100,000 [9].

Decades of studying host-pathogen interactions to understand schistosome infection and disease mechanisms have relied heavily on experimental models of infection [10](); such animal models allow for more experimental control compared to human epidemiological studies. However, despite its public health importance, the mechanistic pathways of urogenital schistosomiasis sequelae are poorly defined, particularly events in the early stages of infection leading to chronic morbidity [11] . This has largely been due to the lack of suitable animal models for *S. haematobium* infection. For instance, infection of mice, hamsters, or other rodents with *S. haematobium* cercariae usually results in involvement of the liver and intestines, rather than the urogenital disease seen in infected humans [12–15]. Non-human primates are known to recapitulate human disease but are difficult to manipulate and are expensive to use as animal models of urogenital schistosomiasis [16].

To address this issue, our group developed a mouse bladder injection model in which *S. haematobium* eggs are directly infused into the bladder walls of mice, allowing us to replicate important pathological features of human urogenital schistosomiasis [17, 18]. Using this model, we demonstrated that a single injection of *S. haematobium* eggs into the bladder walls of mice elicits significant responses in the bladder at both the gross and molecular morphological level. Furthermore, *S. haematobium* egg-induced responses were characterized by regional and systemic Type 2 immune responses, inflammatory cell activation and infiltration, bladder granuloma, bladder fibrosis, egg shedding, hematuria, and urothelial hyperplasia, recapitulating human disease. This has led to subsequent publications [19–26] illustrating the power of this approach.

We sought to leverage this model to determine the early molecular events in the bladder that occur after exposure to *S. haematobium* eggs, and to elucidate the pathways involved in pathology associated with urogenital schistosomiasis. We report here that compared to controls, proteome profiling of *S. haematobium-*infected mouse bladders showed differential expression of proteins important for immune and inflammatory responses, increased protein translation or turnover, oxidative stress responses, reduced epithelial barrier integrity, and carcinogenic pathways. These observations offer mechanistic insight into host-parasite interactions and the myriad pathology of urogenital schistosomiasis and may provide clues for improved therapeutic strategies.

## Methods

### Ethics statement

All animal experiments described in this study were carried out in accordance with relevant U.S. and international guidelines. Experimental protocols were reviewed and approved by the Institutional Animal Care and Use Committee (IACUC) of the Children’s National Hospital, Washington DC, United States (protocol #00030764). These guidelines comply with the U.S. Public Health Service Policy on Humane Care and Use of Laboratory Animals.

### Mice

Fourteen to sixteen-week-old female BALB/c mice (known to exhibit marked pathology from schistosomiasis [18]) were purchased from Jackson Laboratories. All experimental procedures were carried out in accordance with the IACUC protocols, and the institutional guidelines set by the Children’s National Research Institute, Children’s National Hospital. Mice were fed *ad libitum* with a standard mouse chow.

### *S. haematobium* egg isolation and preparation

*S. haematobium*-infected Golden Syrian LVG hamsters were obtained from the National Institutes of Health-National Institute of Allergy and Infectious Diseases (NIH-NIAID) Schistosomiasis Resource Center at the Biomedical Research Institute (Rockville, MD, USA). To retrieve and isolate *Schistosoma* eggs from the liver and intestines, hamsters were sacrificed at 18 weeks post-infection when *Schistosoma* eggs in the liver and intestines were at maximum levels [27]. Livers and intestines were minced and homogenized in a Waring blender, resuspended in 1.2% NaCl containing antibiotic-antimycotic solution (100 units Penicillin, 100 µg/mL Streptomycin and 0.25 µg/mL Amphotericin B, Sigma-Aldrich), passed through a series of stainless-steel sieves with sequentially decreasing pore sizes (450 µm, 180 µm, and 100 µm), and finally retained on a 45 µm sieve. To enrich the population for mature eggs and further clean the eggs, the eggs were placed in a petri dish with cold phosphate-buffered saline (PBS) and swirled several times so that mature eggs accumulated in the center of the dish. The process of swirling and harvesting eggs was repeated several times until all the mature eggs were isolated. These eggs were then immediately resuspended to a concentration of 3,000 eggs in 50 μL PBS and retained on ice. Liver and intestine tissue lysates from age-matched, uninfected LVG hamsters (Charles River Laboratories) was prepared as described for *S. haematobium*-infected LVG hamsters for control injections [18].

### Mouse bladder wall injection and urogenital schistosomiasis model

We have previously described a mouse model of *S. haematobium* egg-induced immunopathology that is based on bladder wall injection with *S. haematobium* eggs [17, 18]. Here, we used this model to recapitulate the spectrum of human urogenital schistosomiasis-associated urologic pathology. In brief, mice were anesthetized with 2% isoflurane, placed on a heating pad, and the abdomen depilated. A 2.5 cm midline abdominal incision was made, and the bladder delivered into the surgical field. A 29-gauge needle attached to a 100 µL glass Hamilton syringe was used to deliver 3,000 *S. haematobium* eggs (in 50 μL) into the bladder wall. Control mice were injected with uninfected hamster liver and intestine tissue extract [18]. We have previously shown that lipopolysaccharide contamination of these preparations, including egg-containing hamster tissues, is minimal [18]. The incision was closed with Vicryl suture followed by application of a topical antibiotic, and mice allowed to recover while being monitored until fully conscious. Mice were administered pain medication (5 mg/kg subcutaneous carprofen) for the first 24 hours as needed, and daily monitoring post-surgery was performed until day 7-post injection, at which time mice were euthanized to harvest their bladders.

### Bladder harvesting and preparation for proteomics analysis

Mice were euthanized by CO_2_ overdose with cervical dislocation, their abdomens opened aseptically, and their bladders harvested. Bladder tissues were rinsed with PBS to remove any residual blood and surrounding connective and fatty tissue was removed aseptically. Tissues were blotted with absorbent paper, placed in cryovials, and snap frozen in liquid nitrogen for 15 minutes. Frozen samples were immediately transferred to a -80 freezer for short-term storage prior to cold-chain shipment to the Beijing Genomics Institute (BGI, San Jose, USA) for further processing and analysis as described below.

### Bladder tissue preparation

Samples were transferred into microcentrifuge tubes containing lysis buffer (9 M urea, 50 mM HEPES (4-(2-hydroxyethyl)-1-piperazineethanesulfonic acid), pH 8) supplemented with protease inhibitor, and vortexed gently. To complete the lysis, samples were homogenized for 1 min and then supersonicated for 30 sec at 20% amplitude (Qsonica, Q500 Sonicator). Samples were spun down with a tabletop centrifuge and the protein concentration was determined using the BCA protein assay kit (Thermo Fisher Scientific, Cat: A532225) according to manufacturer’s instructions. 150 µg of each sample was taken and normalized to the same volume (25 µL) with lysis buffer; 2.5 µL of 100 mM Dithiothreitol (DTT) was added to each sample and incubated at 60LJ C for 15 mins (Eppendorf, Thermomixer C) and alkylated using 2.5 µL of 200 mM iodoacetamide (IAM) at room temperature for 20 mins in the dark. The alkylated samples were then quenched with the same aliquot of 100 mM DTT to eliminate excess IAM in the samples. Water (420 µL) and HEPES (50 µL, 100mM, pH 8.5) were added to each sample so that the 9 M urea concentration was below 2 M for enzymatic digestion. Samples were digested by adding 10 µL of Tryp/LysC (0.15 ug/µL, Thermo Fisher Scientific), and incubated at 37LJC overnight (12 hours) with shaking (Eppendorf, Thermomixer C). An additional 5 µL of same concentration Tryp/LysC was added and samples were incubated for another 4 hours. To quench the trypsin reaction, 65 µL of 10% Trifluoroacetic acid was added to each sample [28]. Samples were cleaned up with 100 mg C18 SPE cartridges (Cat # 60108-302, ThermoFisher Scientific) and elutes from each sample were dried down in a SpeedVac concentrator (Savant^TM^ SpeedVac^TM^ SPD120, ThermoFisher Scientific) for TMT labeling.

### TMT labeling

Each dried sample was resuspended with 40 µL 100 mM HEPES (pH 8.5), gently vortexed and then centrifuged. New Tandem Mass Tag (TMT) labels were resuspended with 80 µL Acetonitrile, (Optima^TM^, LC/MS grade, Fisher Chemical^TM^) and 20 µL of TMT reagent was added to corresponding samples as indicated in **S1 Table**. All samples were incubated at room temperature for 1 hour [29]. A pooled sample consisting of 1 µL of each TMT labelled sample was prepared and used as a quality control for efficacy of the TMT labeling process [30]. The TMT label check sample pool was mixed with 60 µL reconstitution buffer (1% Formic Acid) and injected into the mass spectrometer. LC/MS run of label check sample indicated a label efficiency of about 99%. The TMT labelling reaction was quenched with 15 µL of 5% Hydroxylamine solution. The TMT labelled samples were mixed and dried down with Thermo SpeedVac (Savant^TM^ SpeedVac^TM^ SPD120, ThermoFisher Scientific) for LC-MS analysis (workflow details in **S1 Fig**).

### High pH reverse phase HPLC fractionation

The dried down TMT labeled mixture was reconstituted with 80 µL deionized water and fractionated by High pH reverse phase high performance liquid chromatography (HPLC). (Vanquish HPLC, Thermo Fisher Scientific); a total of 96 fractions were collected in a 75 mins method. Eight fractions were strategically selected and combined into 1 fraction; thus 96 fractions were combined into 12 final fractions. Combined fractions were dried down and desalted using C18 stage tips (Cat # PTR-92-05-18, PhyNexus Inc.). Mobile phase A was deionized water with 20 mM Formic Acetate, pH 9.3, and mobile phase B was Acetonitrile (Optima^TM^, LC/MS grade, Fisher Chemical^TM^) with 20mM Formic Acetate, pH 9.3. Details of the separation gradient is shown in **S2 Table**.

### Liquid chromatography – tandem mass spectrometry (LC-MS)

All fractionated samples were analyzed using a Thermo Orbitrap Mass Spectrometer (Orbitrap Eclipse Tribrid Mass Spectrometer) equipped with a nano flow HPLC (Ultimate 3000, Thermo Fisher Scientific). The Nanospray Flex^TM^ Ion Source (Thermo Fisher Scientific) was equipped with a Column Oven (PRSO-V2, Sonation) to heat up the nano column (PicoFrit, 100 µm x 250 mm x 15 µm tip, New Objective) for peptide separation. A high-field asymmetry ion mobility spectrometry (FAIMSpro) compartment was installed in the front end of the Tribrid Eclipse for improved peptide signal to noise ratio. The nano LC method was water acetonitrile-based, and 150 minutes long with 0.30 µL/min flowrate. For each TMT fraction, all TMT labeled peptides were first engaged on a trap column (Part No. 160454, Thermo Fisher) and then were delivered to the separation nano column by the mobile phase (Gradient information indicated in **S3 Table**). A TMT specific MS/MS-based mass spectrometry method on Orbitrap Eclipse was used to sequence TMT peptides that were eluted from the nano column. For the full MS, 120,000 resolution was used, and the scan range was 300 m/z – 1500 m/z. For the data dependent MS/MS (dd-MS/MS), 60,000 resolution was used. Isolation window was 0.7 Da with a fixed first mass of 110.0 Da. Normalized collision energy was set to 32 with a 15-cycle loop. Both full MS and dd-MS/MS were set to ‘Standard’ for AGC target (i.e., the ion population in the Orbitrap mass analyzer) and ‘Auto’ for max injection time (Max IT).

### Quantitative proteomic analysis

LC-MS data were analyzed by Proteome Discoverer version 2.5 (ThermoFisher Scientific). Because all peptides were labeled with TMT tags, TMT quantitative proteomics searches were performed. All searches were performed in Sequest HT node with mass tolerance of 20 ppm MS [28] and 0.05 Da for MS [29], using the *Mus musculus* database (UP000000589) from Swiss-Prot. Percolator node was used for peptide FDR filtering (Stricted: 001, relaxed: 0.05). TMT labelled peptides abundances were normalized by the total peptide abundance.

### Data processing and analyses

The quantified data was uploaded in Microsoft Excel (version 16), and the R statistical environment (www.bioconductor.org; www.r-project.org) was used for downstream analysis. Prior to any downstream analysis, a data integrity check was performed. To limit the number of false identifications, proteins were considered positively identified when they met the following criteria: False discovery rate (FDR) ≥ 95% (i.e., high=99% or medium = 95% confidence), ≥2 unique peptides, and ≥4 peptide spectrum match (PSM). Overall, protein concentration values should be non-negative and without missing values, these cause difficulties in data normalization and downstream analysis. Hence for analysis purposes, proteins had to meet an additional criterion of being quantified in at least two-thirds of the replicates in a group. Since missing or zero values were caused by proteins with abundance below the detection limit, and not a mere absence, any of such (if present) were replaced by 1/5^th^ the minimum positive value of the corresponding variables for each protein, assumed to be the detection limit.

Overall, 9023 proteins were identified and quantified across all samples. For the 9023 proteins identified, 5988 proteins met the inclusion criteria for further analysis (i.e., 219 proteins with FDR<95%, 1517 proteins with <2 unique peptides, and 1299 proteins with PSM<4 were excluded). After data filtering, two proteins with missing abundance values across all samples were excluded and 5986 proteins were used for further downstream analysis.

An unsupervised Principal Component Analysis (PCA) was employed to assess group separation in the data set, and hierarchical cluster analysis employed to assess clustering trends; hierarchical clustering parameters included a similarity measure– Euclidean distance, and a clustering algorithm – Ward’s linkage (clustering to minimize the sum of squares of any two clusters). PCA analysis was performed using the *prcomp* package and hierarchical clustering was performed with the *hclust* function. For an informative first-hand look at the data set, data were normalized by range scaling (mean-centered and divided by the value range of each variable; **see S1 File**) and an initial PCA and hierarchical cluster analysis was conducted. Based on the initial PCA and cluster analysis, two outlier samples (control group, n=1; cases, n=1) were excluded to improve statistical power (**See S2 File**), and all downstream analysis included eight samples (4 replicates from each group: cases, n=4; controls, n=4). The final analysis data set did not require normalization prior to data analysis after removal of outliers (**See S3 File**).

Statistical analyses using fold change (FC) and a 2-tailed multiple student’s t-test were used to identify proteins that are potentially significant in discriminating between two groups, and to show protein patterns of change under different conditions. A false discovery rate (FDR) threshold of <0.05 [31] and an abundance ratio (i.e., between the two groups) of >1.2 absolute FC was considered significant for differentially expressed proteins (DEP). A Pearson’s pattern correlation analysis with FDR correction (<0.05) was also used to determine linear/periodic trends, and to show protein variation patterns under different conditions.

Proteins were searched against the Gene Ontology (GO) database, for function annotation. GO and pathway enrichment analyses were performed to explore the biological functions and enriched pathways of DEPs between compared groups. The ontology covered three domains: cellular component, molecular function, and biological processes. Protein-Protein Interaction (PPI) network analysis was performed by mapping proteins against the STRING database of known and predicted protein-protein interactions [32]. Protein subcellular localization prediction was performed using the WoLF PSORT bioinformatic tool [33].

### Data access

The mass spectrometry proteomics data have been deposited to the ProteomeXchange Consortium [34] via the PRIDE [35, 36] partner repository with the dataset identifier PXD029852 and at https://doi.org/10.6019/PXD029852

## Results

### Mouse bladder tissue proteome at the host-parasite egg interface

The proteome from the mouse bladder tissue extracts from both groups (n=8, 4 replicates each from case and control groups) were characterized. The top 25 most abundant proteins from the bladder tissue proteome for each group, based on relative abundance, provided clearly distinct profiles between case and control groups (**Fig 1; S4 Table**); the most abundant proteins were unique for each group. The most abundant proteins as identified in the cases included 10 proteins involved in defense responses (accession: P51437, P97457, Q91X17, Q8R2S8, P49290, P11247, O08692, P28293, P11672, Q61646), five involved in cell or biological activity (accession: P15864, P43275, P35174, P43277, P43276), five enzymes or modulators (accession: P13707, Q64444, Q8VCT4, Q9Z2V4, P11034), four structural proteins (accession: P12242, P07744, Q5SX40, Q8CGN5), and one hormone (accession: P01132). The most abundant proteins as identified in the controls included nine structural proteins (accession: Q8BGZ7, Q61765, Q6IMF0, Q9ERE2, Q8BFZ3, P46660, Q61897, O08638-2, O88492), five enzymes or modulators (accession: P63054, P07310, P21550, P54869, Q8R2G4), five involved in cell or biological activity (accession: Q9R1Q8, Q9ER97, P56565, P51576, Q0VBF8), and six proteins involved in defense responses (accession: P01845, P01872, P01592, P06330, P62881, P01867).

**Fig 1.**
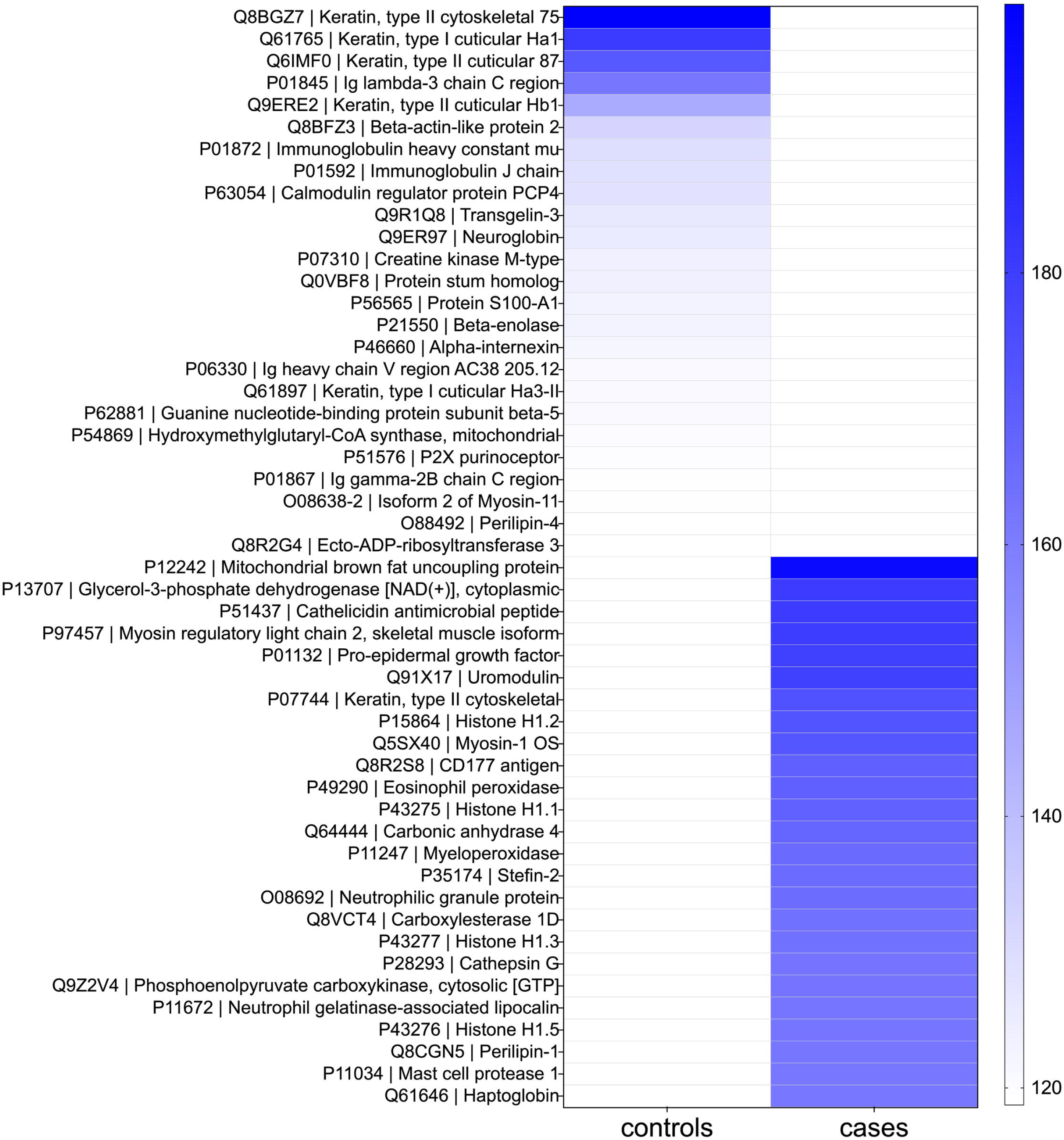
Top 25 most abundant proteins. Heat map shows mean protein abundances from each replicate group. Controls, tissue extract-injection group (n= 4), Cases, egg-injection replicates (n= 4). Proteins are shown as Accession number | Description.

### *S. haematobium* egg-injected bladder tissues show signatures of distinct protein patterns

To characterize the host response to *S. haematobium* eggs in the bladder, we analyzed the data to resolve significant changes in the bladder tissue proteome between the compared groups. PCA and hierarchical clustering was used to initially examine variability and patterns in the data set. PCA showed heterogeneity in components with distinct clustering according to groups and explained 63% of the variation in the dataset across the first two components (**Fig 2a; S2 Fig** (scree plot); **S5 Table**), and replicates within the same group belonged to a single cluster (**Fig 2b**). Fold change (FC) analysis showed that 744 proteins were up-regulated whilst 316 proteins were down-regulated in schistosome infection by >1.2 absolute FC (log_2_FC= ±0.3) (**Fig 2c; S6 and S7 Tables**). Differential protein expression was defined as an adjusted p value (FDR) <0.05 and an absolute fold change >1.2. Based on this criterion, 45 differentially expressed proteins (DEPs) were identified; 25 proteins were significantly upregulated and 20 were significantly downregulated in schistosome infection (FDR<0.05) (**Fig 2d, S8 Table**). Pattern correlation analysis confirmed that these DEPs showed significant strong increasing (n=25 proteins, r >0.7, FDR <0.05) or decreasing patterns with *S. haematobium* egg-injection (n=20 proteins, r >0.7, FDR <0.05) (**Fig 3a, S9 Table**)**. Fig 3b** shows a heatmap of the mean relative abundance of the DEPs showing the specific up and downregulated proteins compared between the two groups.

**Fig 2.**
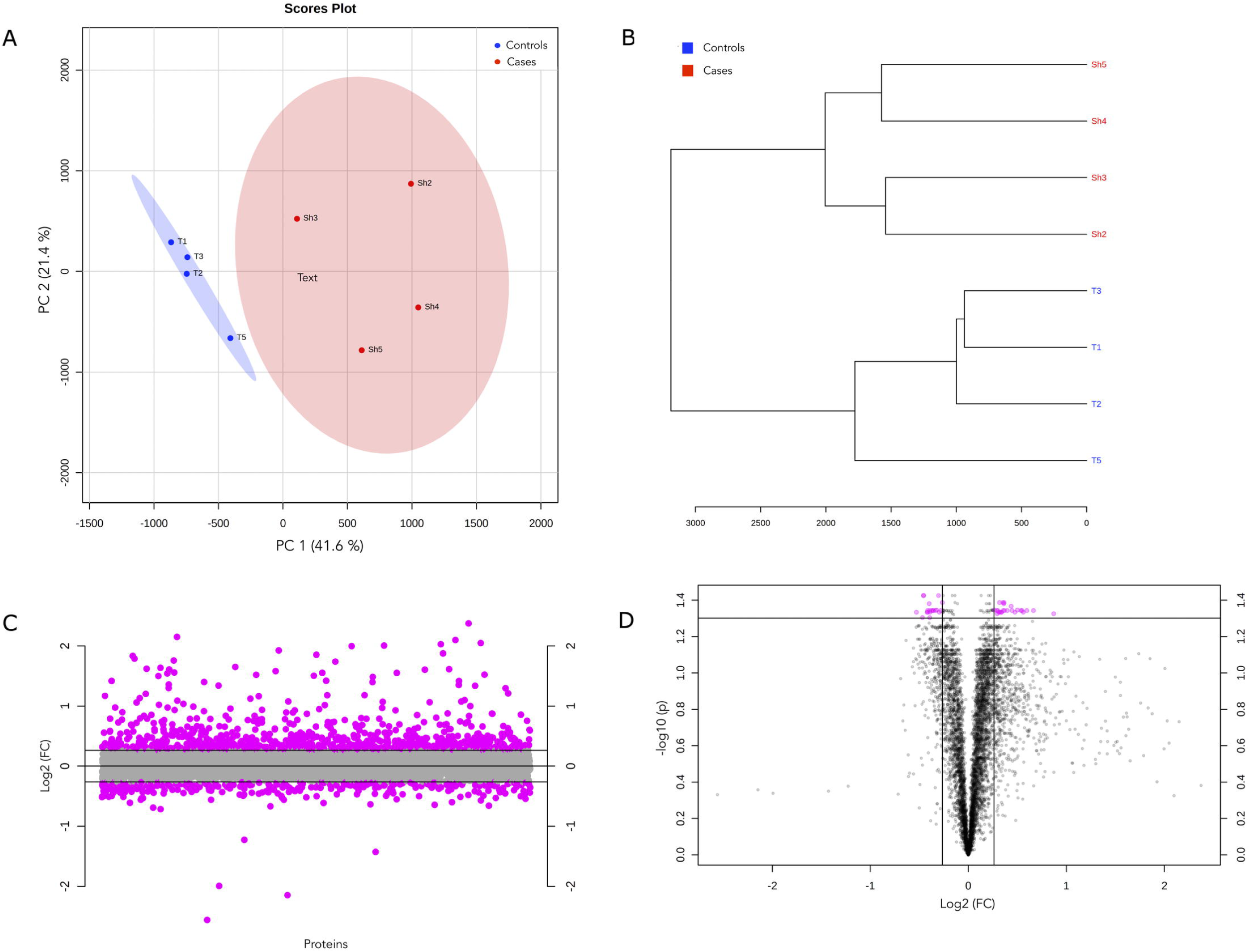
Differential protein expression in the bladder tissue proteome between cases and controls. a) Scores plot from principal component analysis (PCA) across samples, annotated by group. The explained variances are shown in brackets. b) Protein abundance dendrogram. From abundance data, the Euclidean distance was calculated, and samples were clustered based on distances (clustering algorithm – Ward’s linkage). c) Proteins identified by fold change (FC) analysis of case/control ratio with 1.2-FC threshold. Values are on a log scale to show both up-regulated (positive log scale) and down-regulated (negative log scale) proteins symmetrically. Pink symbols represent proteins above the 1.2-FC threshold. d) Volcano plot of differentially expressed proteins (DEPs, n=45) as measured in bladder tissue. Plot is shown comparing protein abundance in egg-injected bladders relative to tissue extract-injected bladders (i.e., cases/controls). Pink symbols represent proteins above the 1.2-FC (log_2_FC= ±0.3) threshold and an FDR <0.05.

**Fig 3.**
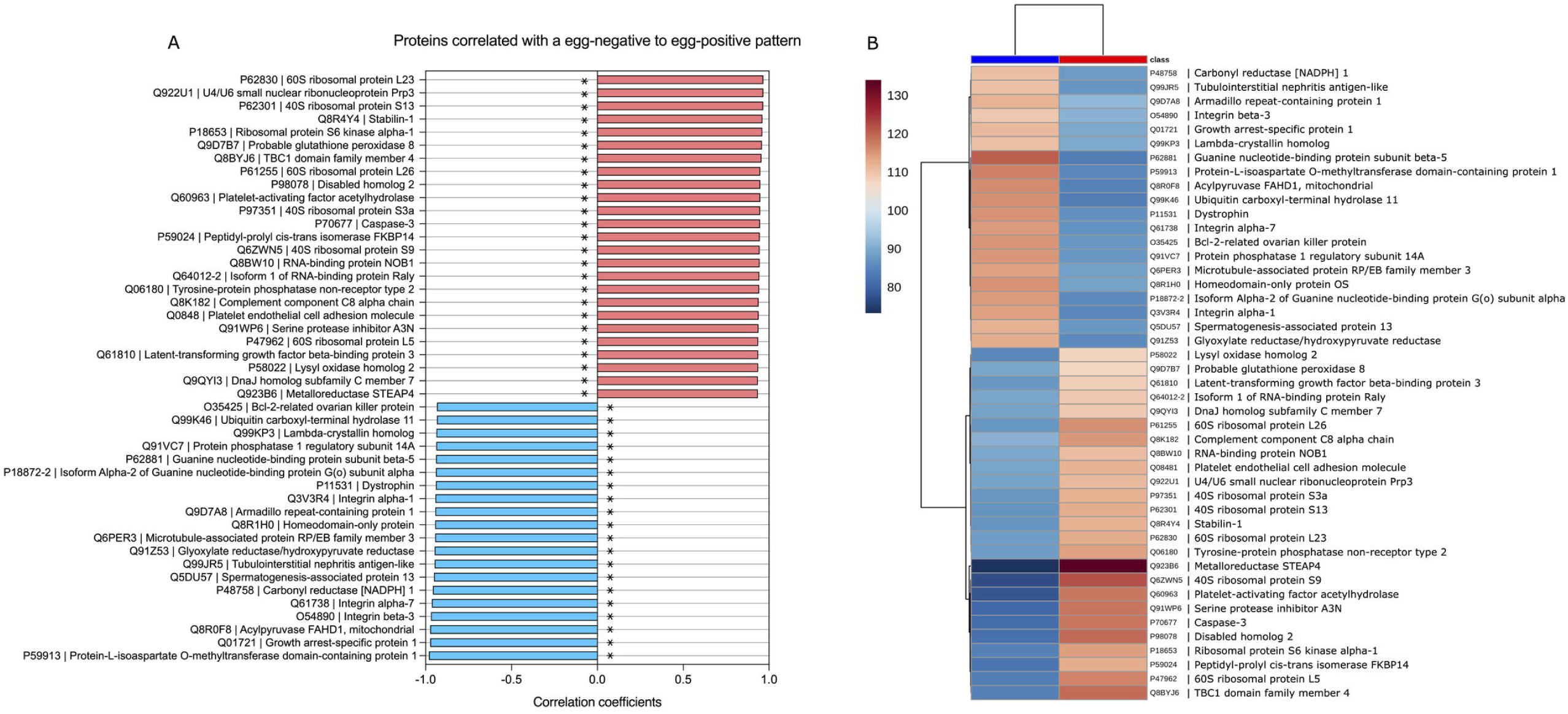
a) Pattern correlation analysis of the 45 proteins, showing increasing and decreasing patterns relative to *S. haematobium* egg status. Correlation coefficients indicate correlation pattern or protein abundance trends relative to the *S. haematobium* egg injection group. Pink and blue bars indicate a positive and negative correlation with *S. haematobium* egg-injection respectively. *, raw p <0.001 and FDR <0.05. b) Clustering and heatmap of significantly differentially expressed proteins among the *S. haematobium* egg-injection group (cases) and the tissue extract-injection group (controls). The heat map was created based on the mean relative abundance of the 45 significant differentially expressed proteins (distance measure using Euclidean, clustering algorithm – Ward’s linkage). Red class label indicates the case group, and the blue class label represents the control group.

### GO enrichment and functional analysis for host responses during *S. haematobium* infection

To infer biological function of the host response to *S. haematobium* egg injection, we used gene ontology (GO) enrichment analysis for the DEPs. Overall, the DEPs were significantly enriched in GO terms predominantly associated with biological processes including cellular processes, metabolic processes, biological regulation, response to stimulus, regulation of biological process, cellular component organization or localization, general multicellular organismal processes, and signaling. These proteins were mostly involved in molecular functions such as binding, catalysis, and regulating molecular and antioxidant activity (**Fig 4a; S10 Table**).

**Fig 4.**
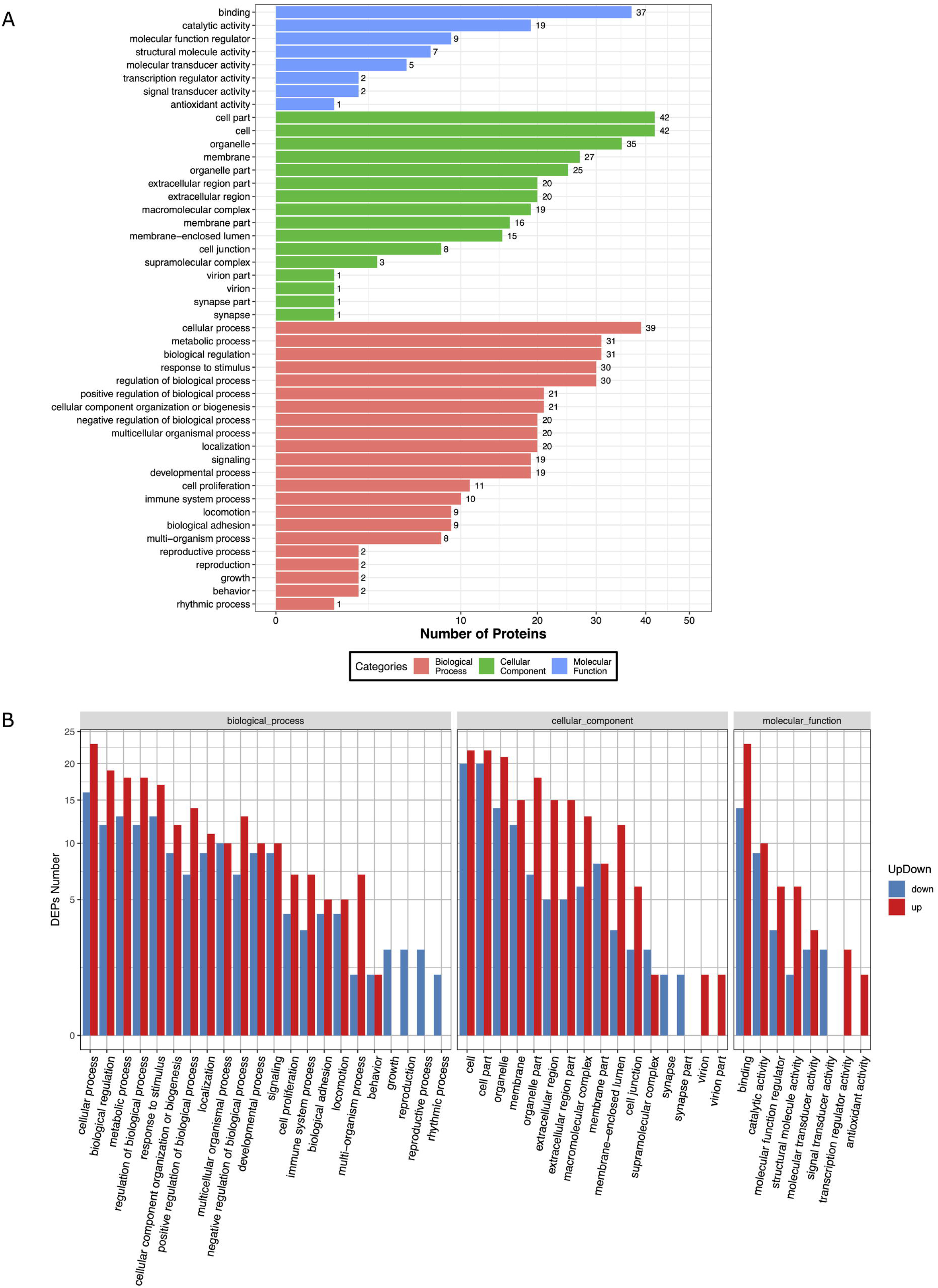
Bar plot showing Gene Ontology Analysis of differential expressed proteins. a) the bar chart shows the distribution of corresponding GO terms. Different colors represent different GO categories. b) the bar chart shows corresponding GO terms based on fold change (FC) analysis of case/control ratio (see **S11 Table** for details).

The proteomic response in the egg injection group was dominated by upregulation of GO terms associated with metabolic processes, response to stimulus, signaling, immune system processes, and a downregulation of growth, reproduction, and rhythmic processes. Molecular functions associated with GO terms such as binding, transcription, antioxidant activity, and regulation of molecular transduction, catalysis, and binding were also upregulated in the egg injection group. (**Fig 4b; S11 Table**). Specifically, these included upregulation of multiple proteins assigned to genes known to be associated with pathways for carcinogenesis (*STEAP4, Pla2g7, Dab2, Fkbp14, Stab1, Loxl2, Prpf3, Ltbp3, Nob1, Raly, Gpx8*), immune and inflammatory responses (*STEAP4, Serpina3n, Dab2, C8a, Pecam1, Prpf3, Dnajc7*), protein translation or turnover (*Rps9, Rpl5, Rpl26, Rps13, Rps3a, Rpl23*), oxidative stress responses (*Pla2g7*), apoptosis (*Casp3*), epithelial barrier integrity (*Ptpn2*), and glucose metabolism (*Tbc1d4*). In addition, downregulated proteins were assigned to multiple genes associated with pathways for carcinogenesis (*Cryl1, Usp11, Gnao1, Gnao1, Grhpr, Ppp1r14a, Hopx*) tumor suppression (*Usp11, Gas1, Cbr1*), cell survival (*Bcl-2*), and reduced structural integrity and cell adhesion (*Itgb3, Itga1, Itga7, Armc1, Usp11, Dmd, Mapre3*).

A protein-protein interaction network, as drawn from the genome, showed that the DEPs formed a highly interactive network, a confirmation that the DEPs identified are at least directly or indirectly related to or belong to the same network; all but 4 of the DEPs (Spermatogenesis-associated protein 13, DnaJ homolog subfamily C member 7, Lambda-crystallin homolog, Carbonyl reductase [NADPH] 1) were interacting within the same network (**Fig 5; S12 Table**). This may also confirm that the identified DEPs taken together, contribute to the complex etiology of pathology and host responses to infection, as well as the characteristic bladder tissue changes in response to infection. Analysis of the subcellular location category showed that many proteins were enriched in the nucleus (26.7%), the extracellular space (24.4%), and cytoskeleton (20.0%) (**S3 Fig 3, S13 Table**).

**Fig 5.**
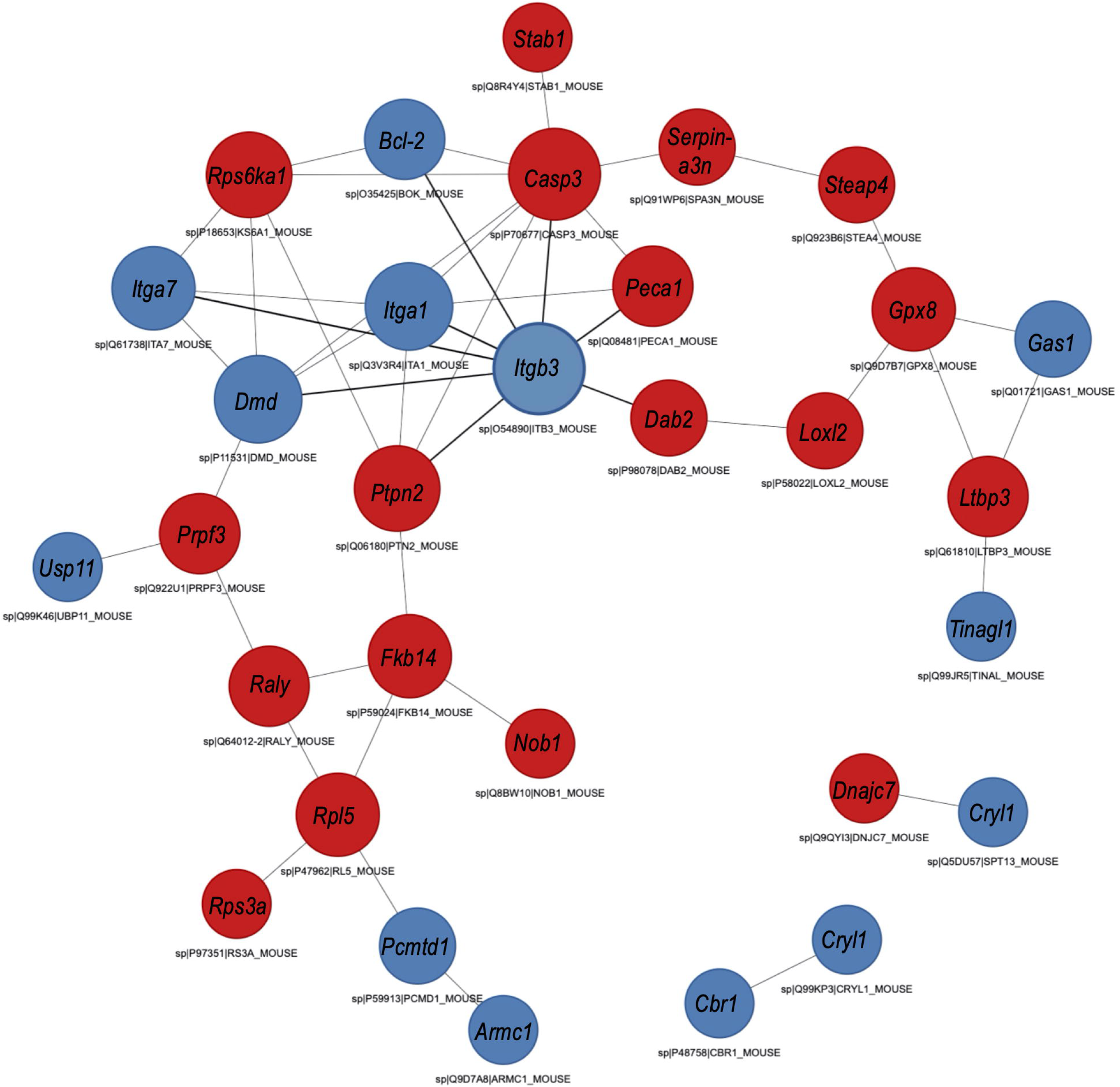
STRING analysis of protein-protein interaction (PPI). Analysis was done considering significant differentially expressed proteins in bladder tissues of mice in the egg-injection group (cases) vs. the tissue extract-injection group (controls). Red nodes represent up-regulated proteins; blue nodes represent down-regulated proteins. Interactions include direct (physical) and indirect (functional) associations from computational prediction, from knowledge transfer between organisms, and from interactions aggregated from primary databases.

Based on our findings, we propose a biological interpretation and model for the proteomic alterations observed and how these may together contribute to the complex dynamics of urothelial pathology and characteristic bladder tissue changes in urogenital schistosomiasis (**Fig 6**). This model demonstrates how the differential expression of the observed proteins contribute to notable pathology including, urothelial hyperplasia, carcinogenesis, active protein translation and enhanced immune and inflammatory responses relevant for wound healing and granuloma formation, induction of Th2 immune responses, and reduced structural integrity important for egg shedding, all of which are characteristic of *Schistosoma* infection. The model is discussed in detail below. Importantly, this model presents a biochemical basis for the histological changes that were observed in a previous study from our group describing the novel mouse model of *S. haematobium* used in the current study [18]. Thus, the changes observed in the urinary bladder proteome in the current study confirms the histological changes observed in our previous study.

**Fig 6.**
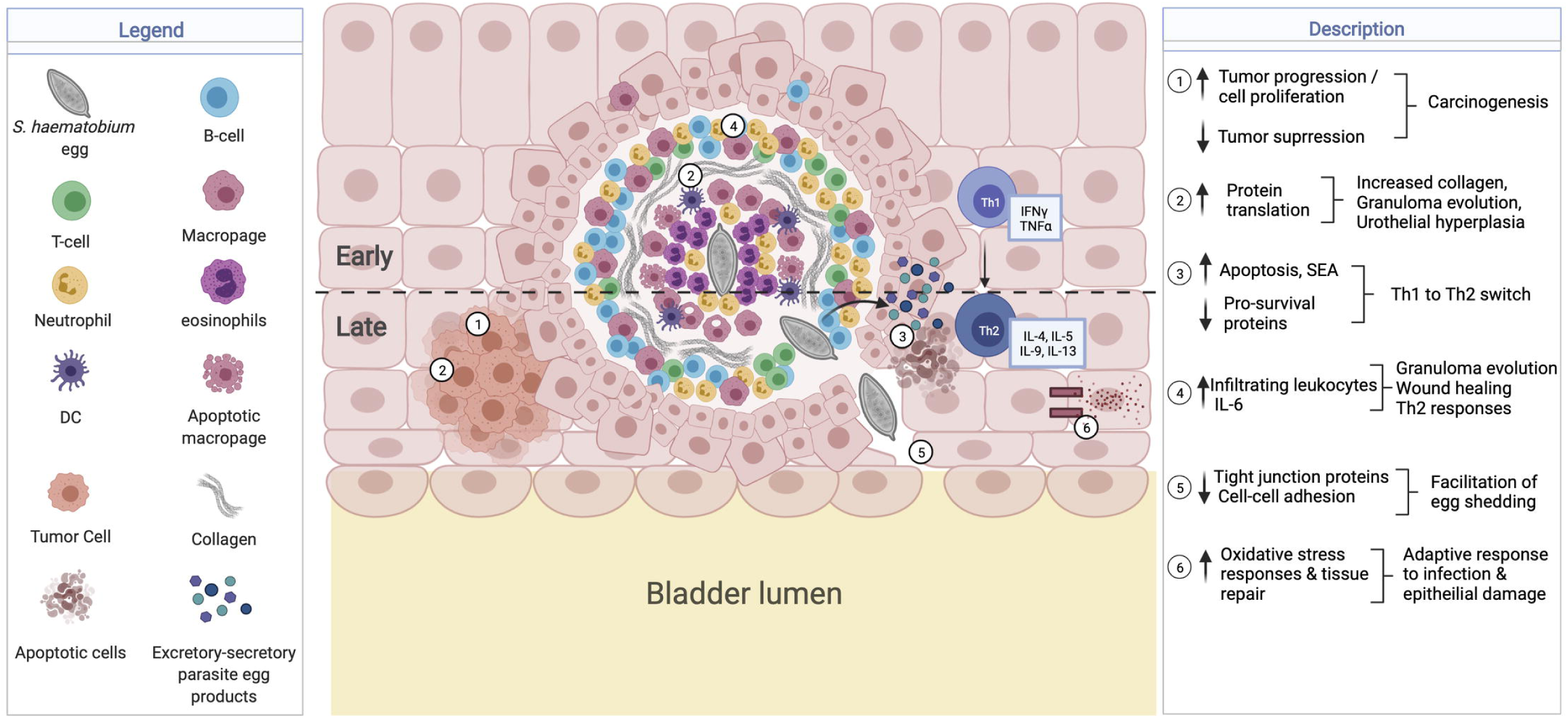
Proposed model for the proteins modified during *S. hae*matobium infections which may contribute to host pathology. Created with BioRender.com ↑ *STEAP4, Pla2g7, Dab2, Fkbp14, Stab1, Loxl2, Prpf3, Ltbp3, Nob1, Raly, Gpx8,* ↓ *Cryl1, Usp11, Gnao1, Grhpr, Ppp1r14a, Hopx, Gas1, Cbr1*, *Bcl-2*) associated with increased cell proliferation and decreased tumor suppression, contributing to carcinogenesis. ↑ *Rps9, Rpl5, Rpl26, Rps13, Rps3a, Rpl23* associated with increased protein translation, collagen transcription and granuloma formation, as well as urothelial hyperplasia. ↑ *Casp3,* ↓ *Bcl-2* in addition to secreted soluble egg antigens (SEA) associated with increased apoptosis and switch in immune responses from Th1 to Th2. ↑ *Dab2, STEAP4, Prpf3, Pecam1* and *Serpina3n* may promote Th2 differentiation, and infiltrating levels of macrophages, neutrophils, dendritic cells, and T-cells, also relevant for granuloma, tissue repair and wound healing. ↓ *Itgb3, Itga1, Itga7, Armc1, Usp11, Dmd, Mapre3* reduce cell-cell adhesions and facilitate egg shedding. ↑ *Pla2g7* and *Ptpn2* may be an adaptive oxidative stress response to promote epithelial barrier integrity.

## Discussion

Schistosome infection remains one of the most important causes of helminth-related morbidity and mortality, with the greatest overall economic and public health impact [1–3]. In urogenital schistosomiasis, most of the morbidity and mortality is mainly due to host immunological reactions to *S. haematobium* eggs deposited within the walls of the urinary tract, with the host bladder undergoing morphological changes at the gross and molecular level. However, the challenges of conducting infection and mechanistic studies in humans have limited our understanding on the mechanisms of the disease [11]. This is compounded by a lack of high-fidelity animal models that can recapitulate human urogenital schistosomiasis.

We have previously demonstrated that injecting *S. haematobium* eggs directly into the bladder walls of mice is capable of recapitulating many features of human urogenital schistosomiasis, including granulomatous inflammation, urothelial hyperplasia, egg shedding, and bladder fibrosis [18]. This approach is based on the premise that schistosome eggs are highly antigenic, and secrete a vast variety of substances including omega-1 [37], kappa-5 [38], interleukin-4-inducing principle of *Schistosoma mansoni* eggs (IPSE)/alpha-1 [39, 40] into host tissues. These substances cause a variety of host responses and influence the surrounding host tissues in schistosomiasis-associated pathogenesis [41]. We used this approach to facilitate identification of perturbed proteins and to elucidate the processes involved in pathology in response to *S. haematobium* eggs. Our proteomic data showed that the most abundantly expressed proteins in the egg-injected group (cases) were mostly involved in defense responses and cell activity, while that of the tissue extract-injected group (controls) were mostly structural and enzyme activity-related proteins. We identified 45 differentially expressed proteins (DEPs); 25 proteins were significantly upregulated and 20 were significantly downregulated in response to *S. haematobium* egg injection. Functional analysis of the DEPs showed that biological processes including carcinogenesis, immune and inflammatory responses, increased protein translation or turnover, oxidative stress responses, reduced cell adhesion and epithelial barrier integrity, and increased glucose metabolism were significantly enriched in *S. haematobium* eggs. Our findings highlight the complexity of the etiology of pathology and host responses to *S. haematobium* infection, and the characteristic bladder tissue changes that lead to pathology and parasite transmission.

Urogenital schistosomiasis is known to predispose individuals to earlier onset and more aggressive bladder cancers, thus infection with *S. haematobium* has been classified as a Group 1 carcinogen by the World Health Organization’s International Agency for Research on Cancer [42, 43]. In the current study, differential expression of multiple proteins implicated in carcinogenesis or cancer-associated pathways were identified, and proteins with roles in cellular activity were amongst the most abundant in the egg-injected group. This is consistent with the observation that mouse bladder wall injection with *S. haematobium* eggs leads to urothelial hyperplasia, a potentially pre-cancerous state [18]. Proteins including metalloreductase (*STEAP4*) [44], platelet-activating factor acetylhydrolase (*Pla2g7*) [45], disabled homolog 2 (*Dab2*) [46], peptidyl-prolyl cis-trans isomerase (*Fkbp14*) [47], stabilin-1 (*Stab1*) [48, 49], lysyl oxidase homolog 2 (*Loxl2*) [50], U4/U6 small nuclear ribonucleoprotein (*Prpf3*) [51], latent-transforming growth factor beta-binding protein (*Ltbp3*) [52], RNA-binding protein (*Nob1*) [53], isoform 1 of RNA-binding protein (*Raly*) [54], and probable glutathione peroxidase 8 (*Gpx8*) [55] are all expressed in many different malignant tissues, including bladder tissue, and are linked with increased cell proliferation, oncogenic properties, and poor cancer prognosis. This is in line with our observations, as all these proteins were more highly expressed in egg-injected mice bladder tissue. Evidence shows that proteins such as *Dab2* play a significant role in aggressive human urothelial carcinoma of the bladder. *Dab2* promotes changes to epithelial-mesenchymal transition and increases tumor proliferation, migration, and invasion; inhibiting the protein leads to suppression of these properties and improves prognosis [46]. Similarly, the downregulation of proteins that play a direct or indirect role in tumor suppression including ubiquitin carboxyl-terminal hydrolase 11 (*Usp11*) [56] [57], growth arrest-specific protein 1 (*Gas1*) [58, 59], and carbonyl reductase (*Cbr1*) [60], lambda-crystallin homolog (*Cryl1*) [61], isoform alpha-2 of guanine nucleotide-binding protein (*Gnao1*) [62], glyoxylate reductase/hydroxypyruvate reductase (*Grhpr*) [63], protein phosphatase 1 regulatory subunit 14A (*Ppp1r14a*) [64], and homeodomain-only protein (*Hopx*) [65] have been linked to cancer-associated pathways in several tissues, including the bladder. The downregulation of *Itgb3* through the action of MIR-320a is also associated with bladder carcinoma invasion in bladder transitional cell carcinomas [66]. The decreased expression of these proteins, as observed in the current study, could contribute to neoplastic changes in the *S. haematobium* egg-exposed bladder. Given that the dysregulation of proteins that control cell proliferation and death represents a core characteristic of cancer, it is not surprising that these proteins were identified as significantly up- or down-regulated in egg-injected mice bladder tissue and highlights the complex bladder morphological changes in response to *S. haematobium* infection leading to pathology. Despite the limited evidence on the role of these proteins in *S. haematobium* studies, these pathways provide an important insight for further investigations to identify host proteins and their role in *Schistosoma*-associated bladder cancer.

Consistent with findings from transcriptional profiling of the bladder in urogenital schistosomiasis [23], we found significant upregulation of several ribosomal proteins, which is an indication of increased protein translation and turnover. Ray and colleagues demonstrated that this may be a response to *S. haematobium* egg deposition in the bladder, where there is high urothelial turnover or hyperplasia, highly active granuloma formation, and increased collagen transcription.

Also relevant to granuloma formation during schistosome infection is apoptosis; soluble egg antigens (SEA) from schistosome eggs have been shown to induce or increase the level of apoptosis [67]. Apoptosis is also associated with the switch in immune responses from type 1 to type 2 during schistosome infection. This is postulated to prevent an exaggerated inflammatory response from egg-induced pathology and to favor parasite survival in the host [68]. The ability to induce apoptosis is thus crucial to schistosome infection, and this is consistent with our observation of downregulation of anti-apoptotic and pro-survival proteins, including the Bcl-2-related ovarian killer protein (*Bcl-2*) and the upregulation of pro-apoptotic proteins such as Caspase-3 (*Casp3*). It is possible that these pathways are stimulated as part of egg-induced pathology and immunomodulatory processes characteristic of schistosome infection [69, 70].

The severe and chronic clinical manifestations of schistosomiasis are immune-mediated, and parasite-specific immune responses cause a downregulation and immuno-modulation of the host’s immune system [69, 70]. Specifically, schistosome-specific immune responses and disease sequelae are mediated by humoral and cellular responses initiated after parasite antigen interaction with antigen-presenting cells, which subsequently direct T-cells towards a T-helper type 1 (Th1), type 2 (Th2), or regulatory T-cell (Treg) phenotype [3, 71]. At the onset of egg deposition, there is an emergence of Th2 responses important for immunomodulation, while Th1 responses are important for protection against infection begin to decrease [69, 71]. Thus, immune responses to schistosome infection is complex and is dependent on a balance between Th1 and Th2 responses as well as antibody-mediated immunity [71]. Bladder wall injection with *S. haematobium* eggs induced an upregulation of several proteins related to granulomatous inflammation and type 2 immunity, with proteins involved in defense responses being the most abundant in the proteome of egg-injected bladder tissues. Proteins such as complement component (*C8a*), platelet endothelial cell adhesion molecule (*Pecam1*), and serine protease inhibitor A3N (*Serpina3n*) were highly expressed owing to inflammatory responses to egg-injection and tissue repair/wound healing [72–74]. The increased expression of metalloreductase proteins in the egg-injected mice bladder tissue may have implications for regulating the production of IL-6 (i.e. an increase) [75], where in parasitic diseases like schistosomiasis, an increase in IL-6 may promote Th2 differentiation and inhibit Th1 polarization [76]. In addition, the upregulation of Disabled homolog 2 (*Dab2*) may facilitate Treg-mediated immunosuppression in response to infection; a downregulation of this protein has been linked with a pro-inflammatory switch [77, 78]. Consistent with granuloma formation, an increase in U4/U6 small nuclear ribonucleoprotein Prp3 (*Prpf3*) has been linked with infiltration of macrophages, neutrophils, dendritic cells, and T-cells [51].

To exit the host epithelium and continue the lifecycle of the parasite, schistosome eggs must cross tight junctions between individual epithelial cells [79]. In experimental mouse studies of *S. haematobium*, Ray and colleagues demonstrated a decrease in transcription of junction adhesion-related genes which occurred in the context of egg shedding [23]. This is consistent with our finding that several proteins involved in structural integrity and cell adhesion, including the integrins (*Itgb3, Itga1, Itga7),* Armadillo repeat-containing protein (*Armc1*), Ubiquitin carboxyl-terminal hydrolase 11 (*Usp11*), Dystrophin (*Dmd*), and Microtubule-associated protein (*Mapre3*) were downregulated in the mouse bladder following egg injection. Whether these proteins are directly or indirectly involved in a compromised epithelium in urogenital schistosomiasis remains undefined. However, given the functional roles of such proteins and the role of schistosome eggs in causing pathology, we speculate that *S. haematobium* eggs induce, or at least exploit, a compromised urothelial barrier that features disrupted cell to cell adhesion (i.e., decreased expression of bladder structural and cell adhesion proteins), to pass into the bladder lumen. In fact, a majority of the most abundant (i.e., upregulated) proteins in the proteome of vehicle-injected bladder tissue were linked to structural integrity and cell adhesion.

There is strong evidence suggesting a correlation between *S. haematobium* infection and increased levels of oxidative stress and tissue repair [80], which is in line with our observation of increased oxidative stress responses in egg-injected bladder tissues. The bladder tissue response to epithelial damage may explain the increased levels of epithelial barrier function proteins such as the tyrosine-protein phosphatase non-receptor type 2 (*Ptpn2*) [79]. In response to invading threats like *S. haematobium* eggs, epithelial cells and resident immune cells communicate with one another to coordinate reinforcement of barrier integrity [79].

While the significant detrimental effects of schistosome infections on host tissue are unarguable, there is evidence of sustained metabolic alterations post-infection, notably, increased glucose metabolism and insulin sensitivity [81]. In fact, while the exact mechanisms of these effects remain to be fully understood, some studies have suggested that such alterations may reduce the occurrence and severity of the metabolic syndrome in infected individuals [81, 82]. In the current study we observed an upregulation of the TBC1 domain family member 4 (*Tbc1d4*), which is known to enhance glucose uptake and insulin sensitivity in tissues [83]. Our findings confirm the role of such proteomic pathways in enhancing host glucose uptake to compensate for the nutrient-poor-conditions in the host, often associated with schistosome infection [84, 85].

In the mouse bladder wall injection model, the complex up- and down-regulation of proteins and their associated pathways (as measured in bladder tissues) arise as a response to the damage associated with the retention and/or traversal of the *S. haematobium* eggs and their products across the urothelium and into the circulation. However, the interpretation of this model has some limitations. Our novel model delivers a single episode of parasite egg deposition and may not entirely reflect the full plethora of the continuous egg deposition by adult worms as with natural human infections. Improvement of the mouse model to facilitate chronic infection with egg-laying adult *S. haematobium* worms would allow for further investigation into the unique mechanistic pathways of *S. haematobium*-induced bladder and systemic pathology. Notwithstanding, our findings provide novel insights into the host responses to the infection, while our egg-injection model remains the only experimentally-tractable model for urogenital schistosomiasis [18].

## Conclusion

To our knowledge, we report here the first characterization of host bladder proteomic changes triggered in response to *S. haematobium* eggs. *S. haematobium* egg deposition in the bladder results in significant changes in proteins and pathways that play a role in pathology. The results obtained from this study provide an in-depth analysis of potential host protein indicators for host-parasite interplay and provide new insights into the complex dynamics of urothelial biology in urogenital schistosomiasis. These include several defense response pathways and characteristic bladder tissue changes that lead to pathology and parasite transmission. The functional role of the proteins identified in this study will need to be explored further in relation to natural human infections.

## Acknowledgements

This work was funded by the GWU Cancer Biology Training Program NIH-T32CA247756 (KI) and NIH-R01DK113504 (MH). *S. haematobium*-infected hamsters and parasite eggs were provided by the NIAID Schistosomiasis Resource Center for distribution through BEI Resources, NIAID, NIH: *Schistosoma haematobium-*exposed Golden Syrian LVG hamsters, NR-21966. We acknowledge Dr. Karl Burgess at the University of Edinburgh for advice and direction with data analysis and interpretation.

## Conflicts of interest

Authors declare that there are no conflicts of interest.

## Author contributions

DNMO and MHH conceived and designed the study. DNMO, KI, and OKL conducted the animal experiments and proteomic study. DNMO, LD, MR and MHH analyzed and interpreted the data. DNMO wrote a draft of the manuscript with the aid of the other authors. All authors were involved in revision of the manuscript. All authors approved the final version of the manuscript for submission.

## Supporting information Captions

**S1 Fig. Experimental workflow.**

**S2 Fig. Scree plot shows the variance explained by principal components from PCA analysis.** The green line on top shows the accumulated variance explained; the blue line underneath shows the variance explained by individual principal components.

**S3 Fig. Differentially expressed proteins categorized by their subcellular localization.** E.R, endoplasmic reticulum; cyto, cytosol; cyto_nucl, cytosol or nucleus; extr, extracellular; extr_plas, extracellular or plasma membrane; mito, mitochondrial; nucl, nucleus; plas, plasma membrane.

**S1 Table. Samples and their corresponding TMT tag channels.**

**S2 Table. High pH reverse phase HPLC fractionation gradient information.**

**S3 Table. nanoLC-MS gradient information.**

**S4 Table. Mean abundances of the top 25 proteins in each replicate group.**

**S5 Table. Principal component analysis (PCA) loading scores.**

**S6 Table. Full univariate analysis output for comparisons between cases and controls.**

**S7 Table. Fold change (FC) analysis output with threshold >1.2 absolute FC.**

**S8 Table. Analysis output for volcano plot for differentially expressed proteins.**

**S9 Table. Correlation pattern analysis output for differentially expressed proteins. Output shows correlation coefficients for proteins with abundance trends relative to the *S. haematobium* egg injection group.**

**S10 Table. Gene ontology (GO) analysis for function annotation of differential expressed proteins.**

**S11 Table. Gene ontology (GO) analysis for function annotation of differential expressed proteins based on fold change (FC) analysis of case/control ratio.**

**S12 Table. Protein-Protein Interaction (PPI) network analysis output for differential expressed proteins.**

**S13 Table. Protein subcellular localization prediction analysis for differential expressed proteins.**

**S1 File. Data distribution and normalization.**

**S2 File. Principal component analysis (PCA) and cluster dendrogram for data distribution and outlier assessment.**

**S3 File. Data distribution after removal of outliers.**

## Notes

https://doi.org/10.6019/PXD029852

### Competing Interest Statement

The authors have declared no competing interest.

