## Supplementary figures and images for "Host tissue proteomics reveal insights into the molecular basis of *Schistosoma haematobium*-induced bladder pathology"

### S1 Fig

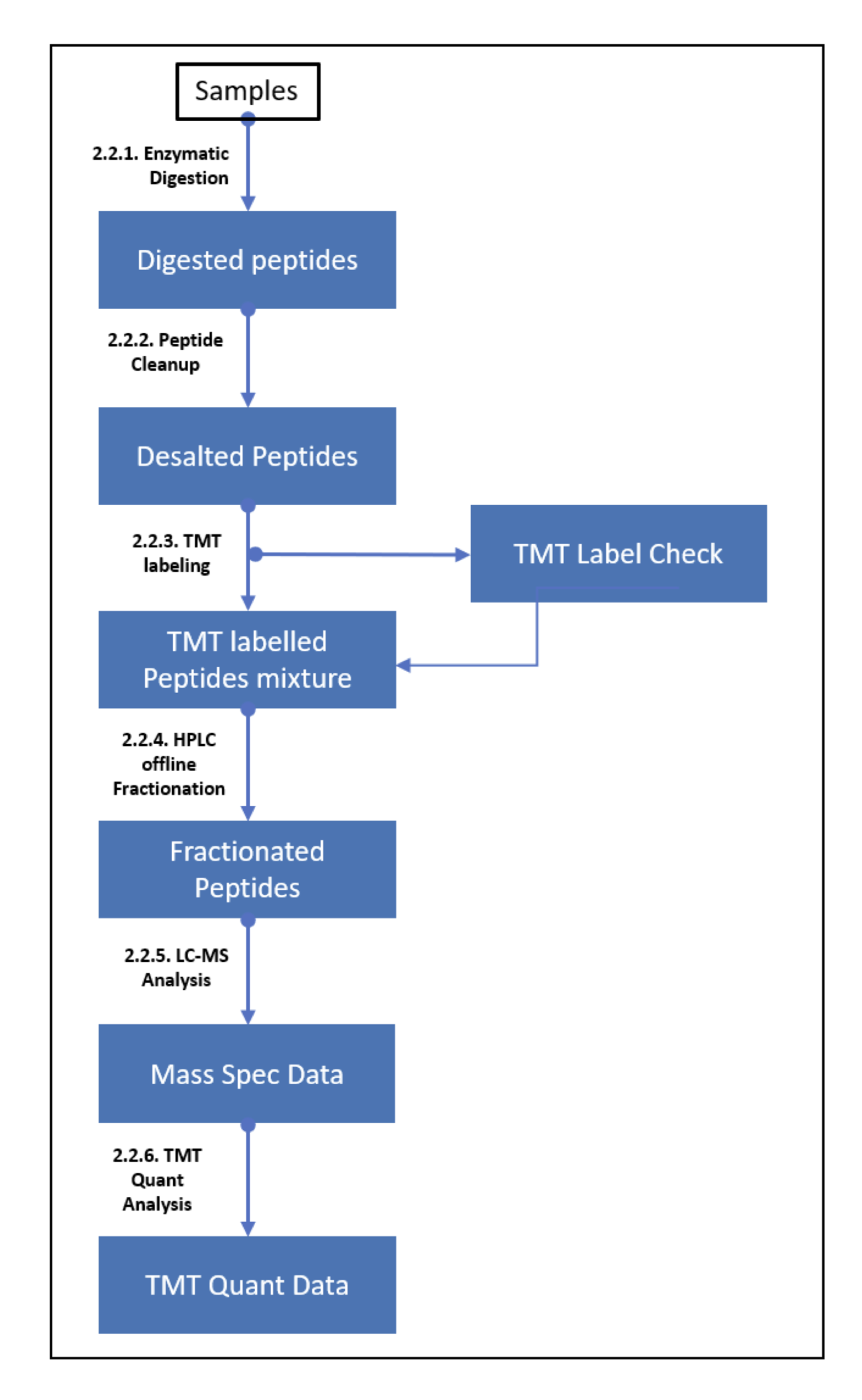

### S2 Fig

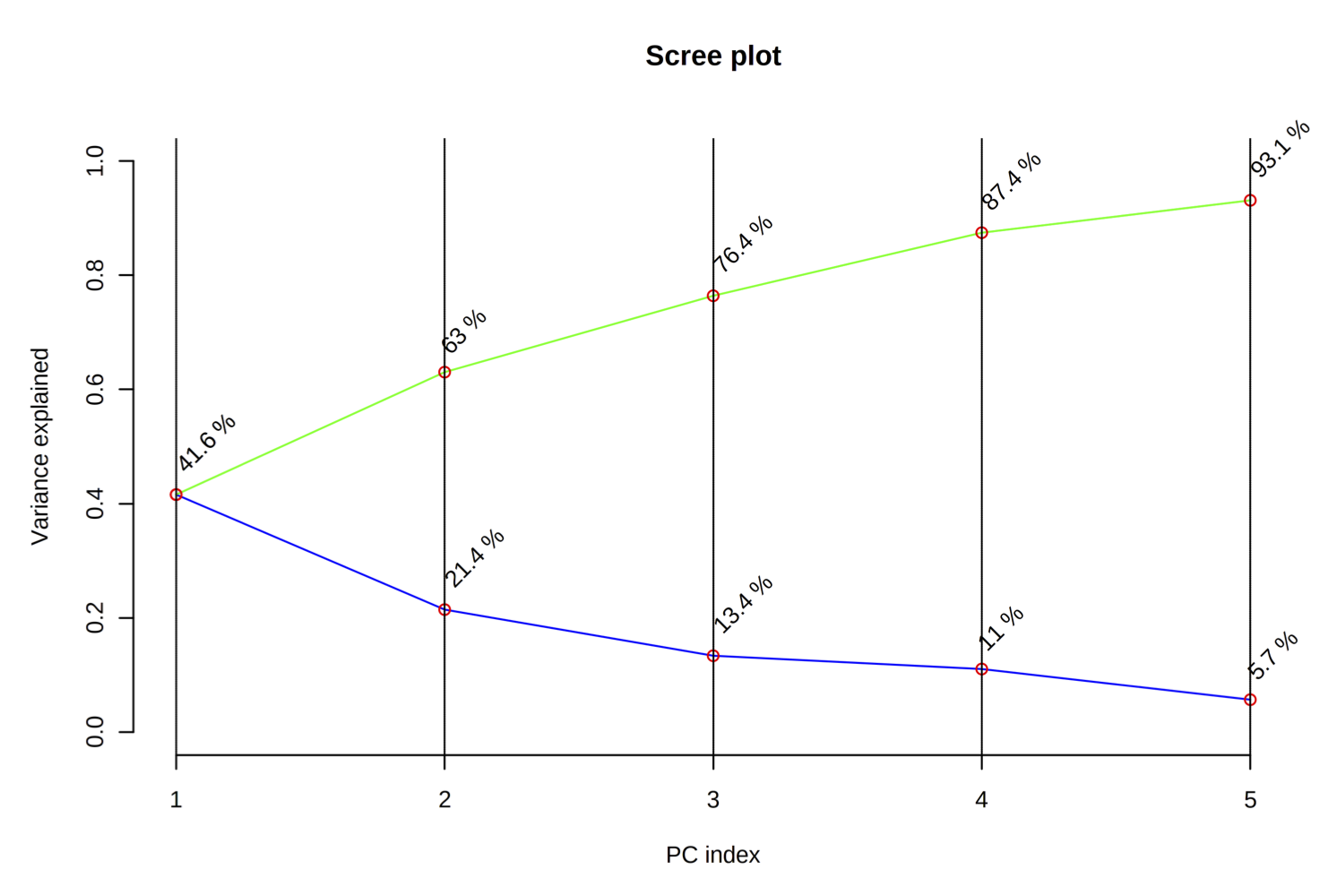

### S3 Fig

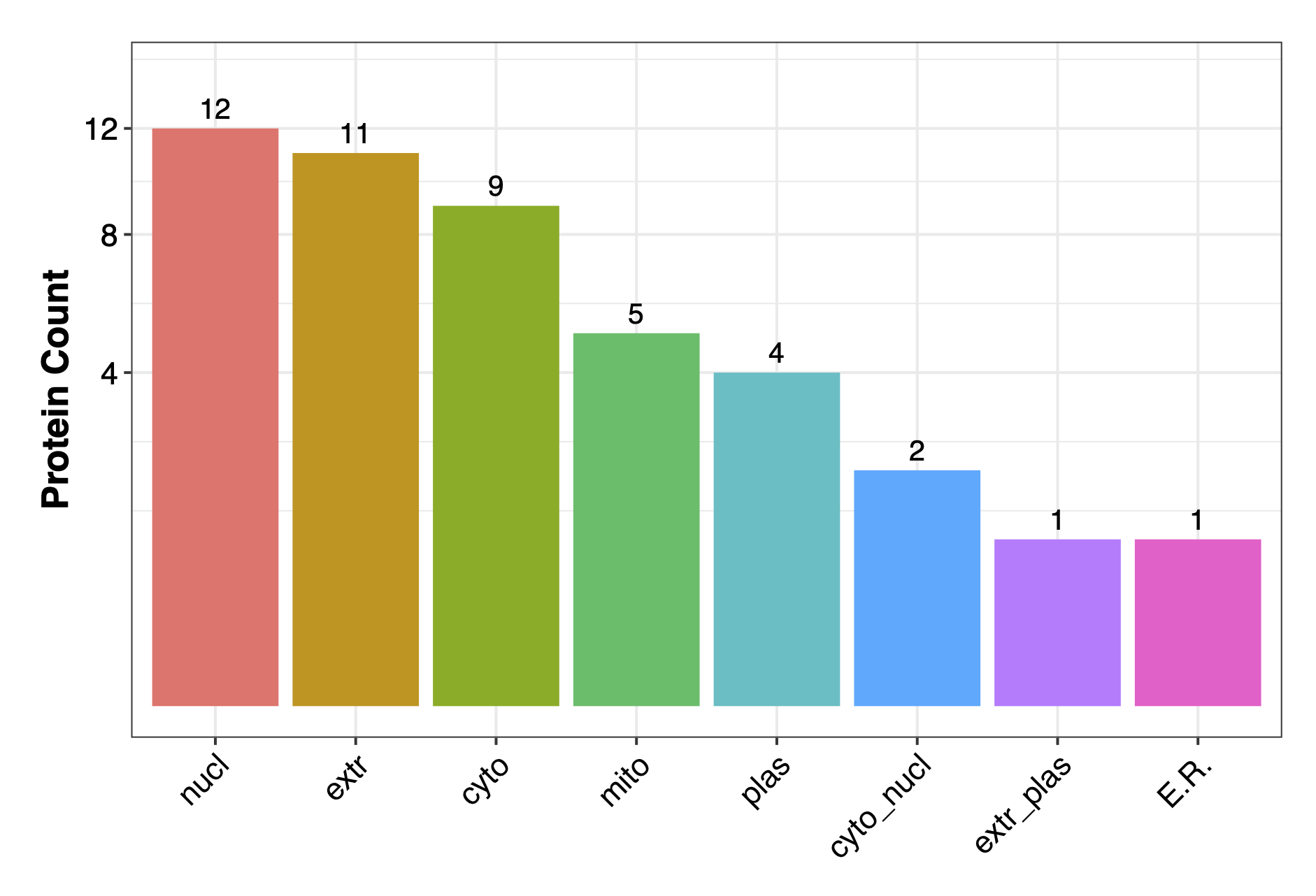
