## Supplementary material for "Host tissue proteomics reveal insights into the molecular basis of *Schistosoma haematobium*-induced bladder pathology": S1 File

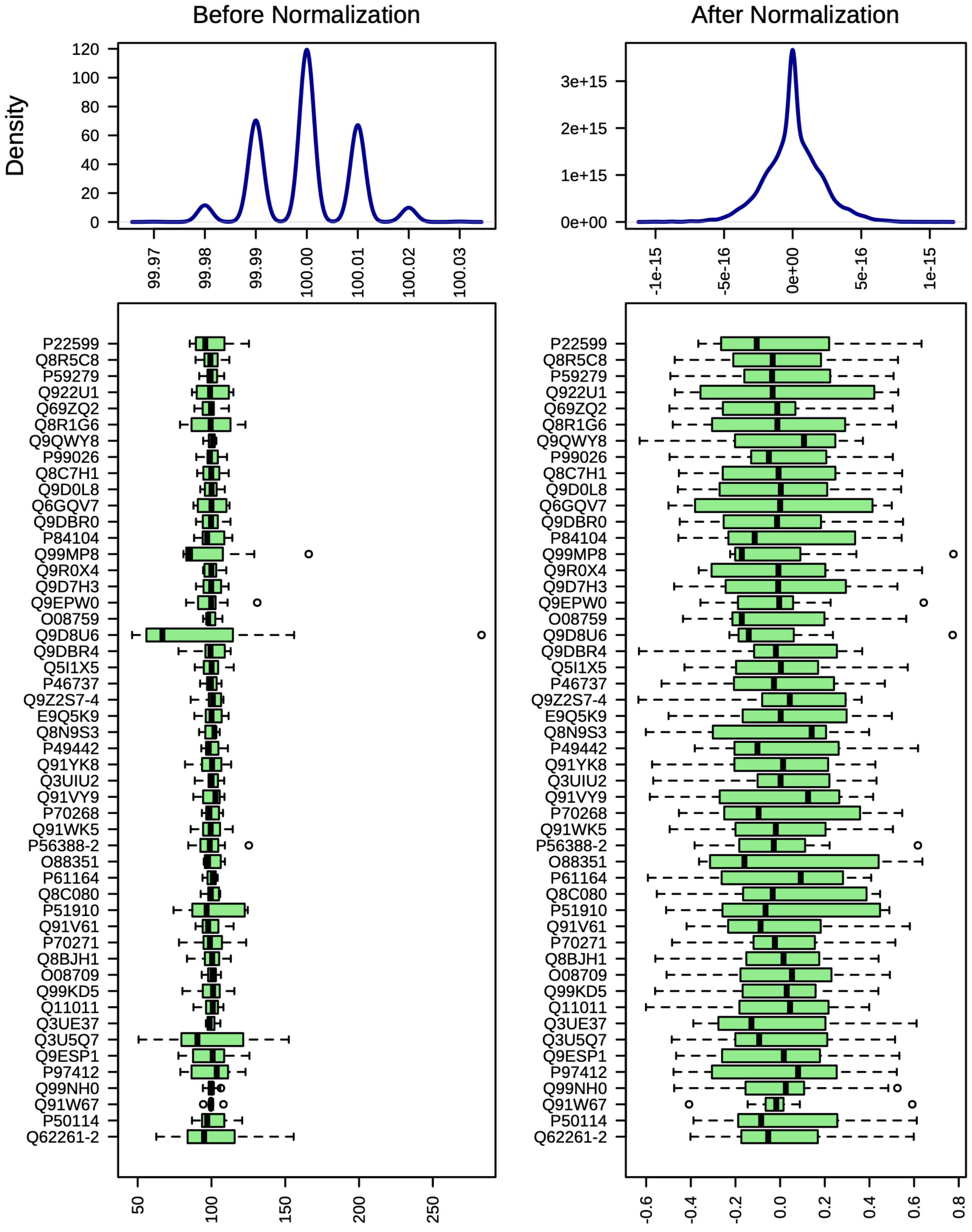


Box plots and kernel density plots before and after normalization (Range scaling) of dataset with all samples (n=10; Cases=5, Controls=5). The boxplots show a maximum of 50 proteins due to space limit. The density plots are based on all samples before and after normalization.
