## Supplementary material for "Host tissue proteomics reveal insights into the molecular basis of *Schistosoma haematobium*-induced bladder pathology": S1 Table

**S1 Table. Samples and their corresponding TMT tag channels.**

| Sample # | Sample Name | Tag Channel |
|---|---|---|
| 1 | T1 | 126 |
| 2 | T2 | 127N |
| 3 | T3 | 127C |
| 4 | T4 | 128N |
| 5 | T5 | 128C |
| 6 | Sh1 | 129N |
| 7 | Sh2 | 129C |
| 8 | Sh3 | 130N |
| 9 | Sh4 | 130C |
| 10 | Sh5 | 131 |
