## Supplementary material for "Host tissue proteomics reveal insights into the molecular basis of *Schistosoma haematobium*-induced bladder pathology": S2 File

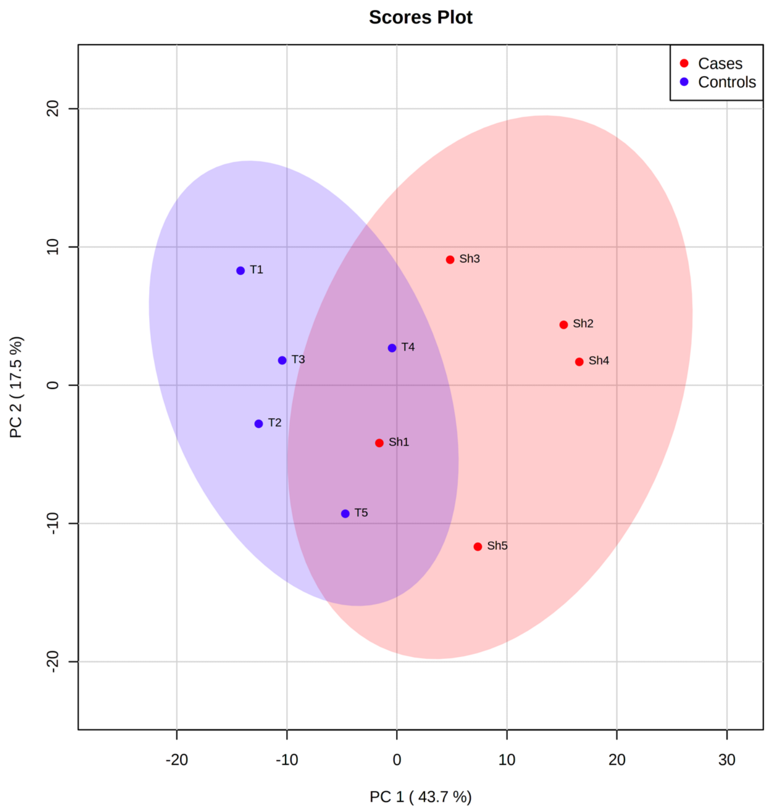


Scores plot from principal component analysis (PCA) across all 10 samples, annotated by group. The explained variances are shown in brackets.

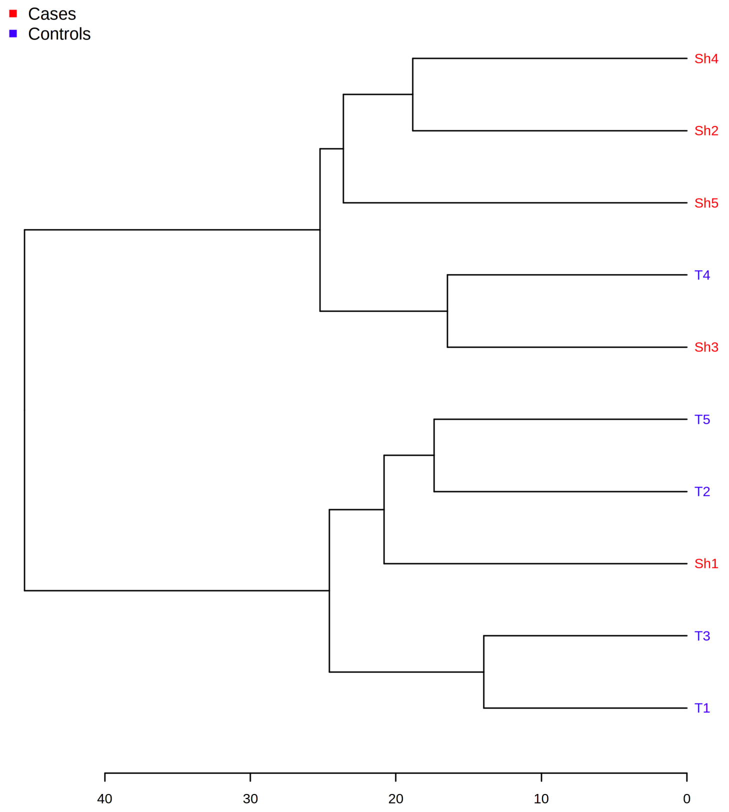


Protein abundance dendrogram for all 10 samples. From abundance data, the Euclidean distance was calculated, and samples were clustered based on distances (clustering algorithm – Ward's linkage). Samples Sh1 and T4 were excluded as outliers.
