## Supplementary material for "Host tissue proteomics reveal insights into the molecular basis of *Schistosoma haematobium*-induced bladder pathology": S2 Table

**S2 Table. High pH reverse phase HPLC fractionation gradient information**

| Time [min] | Flow [ml/min] | %B |
|---|---|---|
| 0.00 | 0.500 | 2.0 |
| 1.00 | 0.500 | 6.0 |
| 12.00 | 0.500 | 20.0 |
| 30.00 | 0.500 | 28.0 |
| 50.00 | 0.500 | 65.0 |
| 53.00 | 0.500 | 98.0 |
| 57.00 | 0.500 | 98.0 |
| 59.00 | 0.500 | 2.0 |
| 60.00 | 0.500 | 2.0 |

*%B, Mobile phase B (Acetonitrile (OptimaTM, LC/MS grade, Fisher ChemicalTM) with 20mM Formic Acetate, pH 9.3).*
