## Supplementary material for "Host tissue proteomics reveal insights into the molecular basis of *Schistosoma haematobium*-induced bladder pathology": S3 File

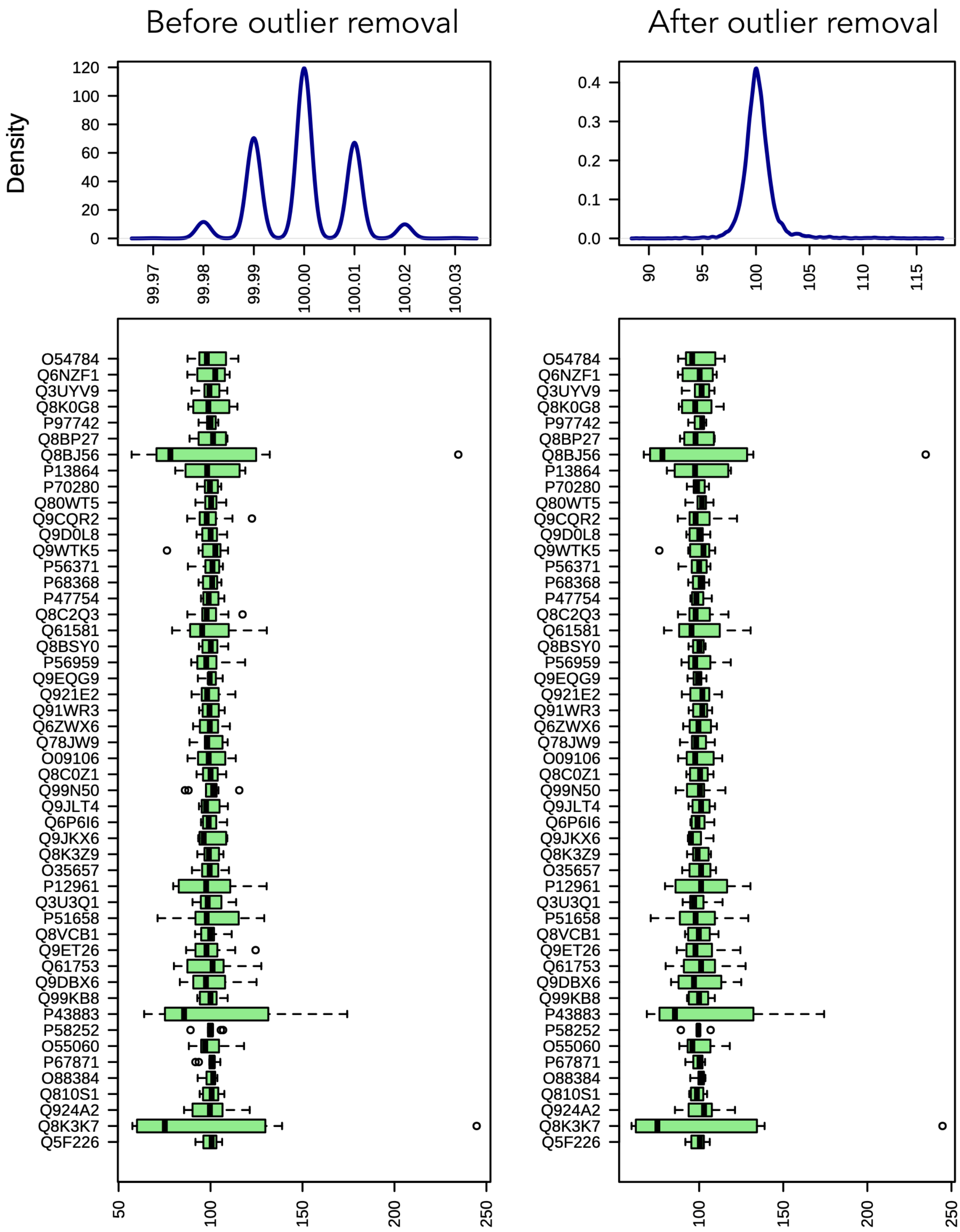


Box plots and kernel density plots before (n=10; Cases=5, Controls=5) and after (n=8; Cases=4, Controls=4) removal of outliers from dataset. The boxplots show a maximum of 50 proteins due to space limit. The density plots are based on all samples before and after outlier removal.
