## Supplementary material for "Host tissue proteomics reveal insights into the molecular basis of *Schistosoma haematobium*-induced bladder pathology": S3 Table

**S3 Table. nanoLC-MS gradient information**

| Time[min] | Flow[µl/min] | %B |
|---|---|---|
| 0.00 | 0.300 | 2.0 |
| 3.00 | 0.300 | 2.0 |
| 3.10 | 0.300 | 2.0 |
| 8.00 | 0.300 | 4.0 |
| 108.00 | 0.300 | 35.0 |
| 128.00 | 0.300 | 65.0 |
| 129.00 | 0.300 | 100.0 |
| 133.00 | 0.300 | 100.0 |
| 134.00 | 0.300 | 2.0 |
| 140.00 | 0.300 | 2.0 |
| 150.00 | 0.300 | 2.0 |

*%B, Mobile phase B (Acetonitrile (OptimaTM, LC/MS grade, Fisher ChemicalTM) with 20mM Formic Acetate, pH 9.3.*
